# Invasive traits of freshwater fish database (ITOFF)

**DOI:** 10.1101/2023.11.15.567195

**Authors:** Aaron Jessop, Anna Michalopoulou, Carolyn Coonan, Lewis Mazzei, Eilidh Sutherland O'Brien, Grace Brady, Carly Davison, Wayne Gourlay, Ethan Henderson, Anna Lornie, Emily McCloskey, Hannah Ramsay, Sophie Wilson, Hideyasu Shimadzu, Miguel Barbosa

## Abstract

**AIM:** Species invasions are a major driver of global biodiversity loss, but only a minority of invasions are successful. Evidence suggests that invasive success is linked to life-history traits. Yet, data on invasive success and species’ traits remain fragmented across multiple sources. Here we present the Invasive Traits of Freshwater Fish (ITOFF) database, an interdisciplinary framework that integrates multiple datasets to elucidate the role of life-history traits in shaping invasive success. ITOFF allows seamless access to invasive species data and fosters collaborative actions through knowledge sharing. ITOFF is supported by an innovative web-application that makes complex relationships between invasive and native species accessible to a broad audience. The scientific contribution of ITOFF is illustrated by examining the role of life-history traits and phylogeny in invasion success.

**LOCATION:** Global.

**METHODS:** Generalized linear models were used to test the contribution of generation time, trophic level, longevity, and temperature range to invasive success. Through divisive cluster analysis we investigate the role of multiple traits in determining invasive success. Finally, we construct phylogenetic trees to investigate the role of evolutionary history in the invasion process.

**RESULTS:** ITOFF unifies data for 1917 freshwater fish species representative of invasive species, those species they endanger, and species impacted by invasives but not considered endangered. Invasive species are generally characterized by greater temperature ranges, but are indistinguishable from impacted, endangered, and critically endangered species for the remaining life-history traits. Further, we show that invasive species are generally not distinct from impacted or endangered species when considering multiple traits or phylogeny.

**MAIN CONCLUSIONS:** ITOFF provides an accessible platform for the improved forecasting of species invasions. ITOFF data shows that classical predictions of life-history traits determining invasive success do not hold amongst freshwater fish species. Forecasting of invasive species must therefore shift towards a wholistic approach encompassing the species and the environment.

## INTRODUCTION

Invasive species are a leading threat to global biodiversity (Reid *et al*., 2019; WWF, 2020). Further, the economic and social costs of invasive species are severe (Andersen *et al*., 2004; Pimentel, Zuniga and Morrison, 2004; Hoffmann and Broadhurst, 2016). Nevertheless, measures can be implemented to mitigate the negative impacts of invasion (Olson, 2006). For example, recreational and commercial fisheries have been used to control invasive lionfish (*Pterois miles*) populations (Ulman *et al*., 2021). However, the control of invasive species can be expensive (Heikkila and Peltola, 2004) and the impacts of invasion are often irreversible (Andersen *et al*., 2004; Heikkila and Peltola, 2004; Keith and Spring, 2013). Prevention is, therefore, the optimal management strategy (Andersen *et al*., 2004). As invasive propensity differs amongst species (Manchester and Bullock, 2000; Sakai *et al*., 2001), species with greater invasive propensity must be identified for the improved efficacy of preventative management (Sakai *et al*., 2001). Life-history traits, such as fecundity and age of sexual maturation, are generally acknowledged to be critical determinants of invasive success (Holway and Suarez, 1999; Deacon, Ramnarine and Magurran, 2011; Chapple, Simmonds and Wong, 2012). However, few comparative studies have identified traits that promote invasive success in freshwater fish species at the global or major geographical region scale (Bernery, 2022). To address this gap, we present the Invasive Traits of Freshwater Fish (ITOFF) database. The ITOFF database provides easy access to research-ready data concerning the life-history traits and impacts of invasive freshwater fish species at the global scale.

Life-history traits that promote invasive success (‘invasive traits’) are predicted to be shared by a significant proportion of successful invasive species (Van Kleunen, Weber and Fischer, 2010). The comparison of trait values between invasive and ‘non-invasive’ species allows the identification of invasive traits (Capellini *et al*., 2015). However, identifying ‘non-invasive’ species has proved challenging because it is extremely difficult to predict if a currently non-invasive species will become invasive in the future (Garcia-Berthou, 2007). For mammalian species, comparing successful and unsuccessful invaders enables the identification of traits critical to invasive success (Capellini *et al*., 2015). This is possible because of the large number of documented, failed mammalian invasions (Capellini *et al*., 2015). However, similar analyses are not appropriate for freshwater fish as introductions are not commonly reported in sufficient detail (Garcia-Berthou, 2007). To circumvent this issue, data can be collated on traits of species that are under threat of extinction from invasive species, because endangered species are unlikely to become invasive themselves. Further, we can incorporate trait data for species known to be impacted by invasives but not considered to be at risk of extinction (here onwards referred to as impacted species). This would be an appropriate control group. Data on the life-history traits of invasive species, those species they endanger, and impacted species have already been recorded (ISSG, 2000; Froese and Pauly, 2022; IUCN, 2022). Nevertheless, these data are stored in fragmented datasets managed by different bodies. Moreover, in their current format, data are not easily accessible for research purposes. The unified and public provision of life-history trait data for invasive species in an immediately accessible and re-usable format for research is urgently needed for the improved, predictive understanding of species invasions.

The ITOFF database creates the unique opportunity to address critical questions related to invasion biology by collating publicly available data on the life-history traits of the most common invasive, critically endangered and impacted freshwater fish species (freely available at https://itoff-dataset.wp.st-andrews.ac.uk). The accessibility of ITOFF is further supported by a web application (available at: https://itoff-dataset.wp.st-andrews.ac.uk/webapplication/) that facilitates the visualization of links between invasive, impacted, endangered and critically endangered species. Networks can be constructed based on the user’s focal species and climate region of choice. To our knowledge, this is the first visualisation of the network of impacts associated with invasive species. These networks reveal the complexity of impacts associated with invasive species. The harmonious incorporation of invasive, impacted, endangered, and critically endangered freshwater fish species in one dataset provides a novel approach to the study of invasion biology, while creating a powerful yet simple tool for future research.

To illustrate the scientific significance and accessibility of ITOFF, we investigate differences in life-history traits often predicted to facilitate species invasions among invasive, critically endangered, endangered, and impacted species for temperate, subtropical, and tropical regions. We focus on four traits shown to be critical to invasive success (Pimm, 1991; Crowder and Snyder, 2010; Zerebecki and Sorte, 2011; Deacon and Magurran, 2016; Rosenthal *et al*., 2021). Namely, generation time, trophic level, longevity, and temperature range. We expected invasive species to have shorter generation times and reduced longevity to facilitate fast population growth (Pimm, 1991) and rapid adaptation (Rosenthal *et al*., 2021). Further, we expected invasive species to have wider temperature ranges, as this increases the likelihood of survival across a range of environments (Zerebecki and Sorte, 2011). Finally, we expected invasive species to be characterised by intermediate trophic levels (i.e. generalists). Generalist species have broader resource use which increases competitive advantage (Crowder and Snyder, 2010) and facilitates behavioural flexibility that in turn promotes survival under novel conditions (Deacon and Magurran, 2016).

There is the possibility that invasive success is not shaped by individual traits but rather by a combination of several. For example, invasive gammarid amphipods cannot be identified from individual traits but can be predicted using ecological profiles that combine multiple characteristics (Grabowski, Bacela and Konopacka, 2007). We therefore test the hypothesis that invasive propensity is not tied to individual traits but arises from a combination of multiple traits. As phylogenetic history influences trait similarity, we additionally utilise the detailed taxonomic information provided by the ITOFF database, to untangle the role of adaptation, chance, and evolutionary history in the invasion process.

## METHODS

### The ITOFF Database

A dedicated worksheet was created for the collection of all data. Data fields include group (i.e., invasive, endangered, or impacted), taxonomy, climate region (divided broadly into temperate, tropical, subtropical, polar, and boreal), IUCN status (a detailed measure of extinction risk), impacts on other species, and life-history traits predicted to be critical to invasive success (Table A1 Appendix).

A list of non-native freshwater fish species was generated using data from the Global Invasive Species Database (GISD; ISSG, 2000), the U.S. Geological Survey’s Nonindigenous Aquatic Species (NAS) database (USGS, 2022), CABI’s Invasive Species Compendium (ISC; CABI, 2022), the R interface to FishBase (Froese and Pauly, 2022), and data provided by the Global Register of Introduced and Invasive Species (ISSG, 2021). The IUCN Red List of Threatened Species (2022) was used to generate three lists: (1) critically endangered and (2) endangered species for which invasive species are considered a threat, and (3) least concern species that are known to be impacted by invasives. Finally, through the screening of invasive species’ profiles, provided in text format by online databases (ISSG, 2000; CABI, 2022; USGS, 2022; Froese and Pauly, 2022), a list of species known to be impacted by invasive species was generated. Cross-referencing this list with the IUCN Red List allowed for the identification of impacted species considered to be of ‘least concern’ in terms of extinction risk; these species were included in the impacted group of the dataset. Scientific names for all species were updated to be in accordance with FishBase (Froese and Pauly, 2022). Further, the database was curated to identify and omit cryptic duplicate entries.

All species in the ITOFF database are categorised as non-native (n = 979), endangered (n = 299), critically endangered (n = 213), or impacted (n = 381). Non-native species are further categorised as invasive, established, extirpated, failed, or reported. Categorisation of non-native species within ITOFF reflects the policy of governments, NGOs, and international agencies by defining species as invasive only if there is an associated negative impact (Rejmanek *et al*., 2002). Species are considered reported if known to have been introduced to a novel ecosystem, but survival beyond introduction is unknown. Reported is the default categorisation for all non-native species. Non-native species that have successfully founded one or more self-sustaining populations are considered ‘established’. Non-native species that have never survived to establishment are considered ‘failed’. Established species that were later eradicated across their non-native range by human management are considered ‘extirpated’. Finally, species are categorised as ‘invasive’ if there are known (or a reported risk of) negative economic, ecological or social impacts associated with the species within its non-native range. It should be noted that of the 535 extant invasive and established species in the ITOFF database, for which IUCN status is reported, only 7 % are endangered or critically endangered.

The ITOFF dataset reports IUCN status and native/non-native status as separate variables. Accordingly, critically endangered, endangered, and impacted species with known non-native populations have been identified and recategorized as non-native within the ITOFF database. Further, impacted species reported by the IUCN Red List of Threatened Species to have undergone large-scale population declines or localised extinctions due to invasive species were recategorized as ‘locally threatened’. This was necessary to ensure that impacted species in the dataset are representative of native species that are tolerant of invaders.

All data were collected by a small team of researchers (N = 11). All researchers followed the same systematic methodology for data collection as outlined below. For all species, online databases, namely FishBase (Froese and Pauly, 2022) and the IUCN Red List of Threatened Species (IUCN, 2022), were screened for data on relevant traits. FishBase was further used for its unique estimate of trophic level, and the life-history tool was used to produce appropriate estimates for key traits (age of sexual maturity, generation time, and longevity) when data were otherwise unavailable. Note that estimates for trait values were only included if data were available for the species or the species’ family. Estimated and observed values for the same trait are recorded in separate columns within the ITOFF database for clarity. For all non-native species, data on traits were additionally collected from species profiles provided by the GISD (ISSG, 2000), ISC (CABI, 2022), and the NAS database (USGS, 2022). Means of introduction was extracted from non-native species’ profiles and the R interface to FishBase (USGS, 2022; CABI, 2022; Froese and Pauly, 2022). Links between species (e.g., predation of an invader on one or more native species) were recorded from species profiles provided by the IUCN Red List of Threatened Species, and above-mentioned databases for non-native species. Links between species were only included if reported to the species level (i.e., reports of an impact by an invasive species on a taxonomic rank higher than the species level are not included in the ITOFF database).

For all traits, if different data sources provided contrasting values, the mean value was used to populate the dataset. Detailed taxonomic information for all species was extracted from the GISD (ISSG, 2000), NAS database (USGS, 2022), and the IUCN Red List of Threatened Species (IUCN, 2021).

To validate the data, hybrids and species of taxonomic uncertainty (i.e., disputed subspecies or inconsistent and unclear nomenclature used in literature) were identified and omitted from the species pool. Conditional formatting rules, created via the Excel Visual Basic for Applications (VBA) tool, were used to control for data entry errors. Namely, entries not matching specified entry formats were visually highlighted, easily recognised, and corrected. Checks were then made in R (R Core Team, 2021) using base functions to ensure consistent formatting was used throughout and to control for duplicates and typos.

### Linking life-history traits and invasive success

We used the ITOFF database to examine differences in life-history traits amongst invasive, critically endangered, endangered, and impacted species in temperate, subtropical, and tropical regions. Polar and boreal regions were excluded from the investigation because of limited data availability. The four traits of interest, generation time, longevity, temperature range, and trophic level, have been systematically reported as key traits for invasive success (Pimm, 1991; Crowder and Snyder, 2010; Zerebecki and Sorte, 2011; Deacon and Magurran, 2016; Rosenthal *et al*., 2021).

Values of life-history traits were compared between the four treatment groups using four generalized linear models (one per trait). Both observed and estimated trait values as reported within ITOFF were included in the analyses. Each full model included trait value as the response variable and species’ status (i.e. invasive, impacted, endangered) as a fixed factor. Separate analyses were conducted for each climate region. Species that occupy multiple climate regions were represented in multiple counts. Locally threatened species were excluded from the analyses as they are representative of native species that have been severely impacted by invasive species within part or all of their sympatric range. Error distributions were chosen based on data type and distribution of the response variable.

### Multiple traits in determining invasive success

To investigate the role of multiple traits in determining invasive success a hierarchical cluster analysis, divisive analysis using the diana function of the Cluster package in R v 4.2.0. (Maechler *et al*., 2022), based on age of sexual maturity, trophic level, and preferred temperature range was performed. We utilised a random subset of 75 tropical species included in the ITOFF database (25 invasive, 25 endangered, and 25 impacted). A subset was used because data on the three traits of interest were often not reported collectively for all species, and data availability differed between treatment groups. To investigate the relationship between adaptation and evolutionary history in the context of invasive species, we constructed a taxonomic tree utilising the species pool used for the cluster analysis.

## RESULTS

### ITOFF Database

Data availability and sample size differ amongst species categories within ITOFF. Limited data availability reflects knowledge gaps which can be mitigated with the ITOFF database. For example, of all non-native species in the ITOFF database, only 31 are recorded in polar climatic zones. Further, relatively few species are reported as failed invaders, and data availability within this group is limited.

The ITOFF web application, clearly shows that endangered and impacted species are affected by multiple invasive species (Figure 1). Further, invasive species are shown to have impacts on other invasive species. Information on the type of impact (i.e., competition, predation), and profiles for the species selected are provided when using the web application directly.

**Figure 1.**
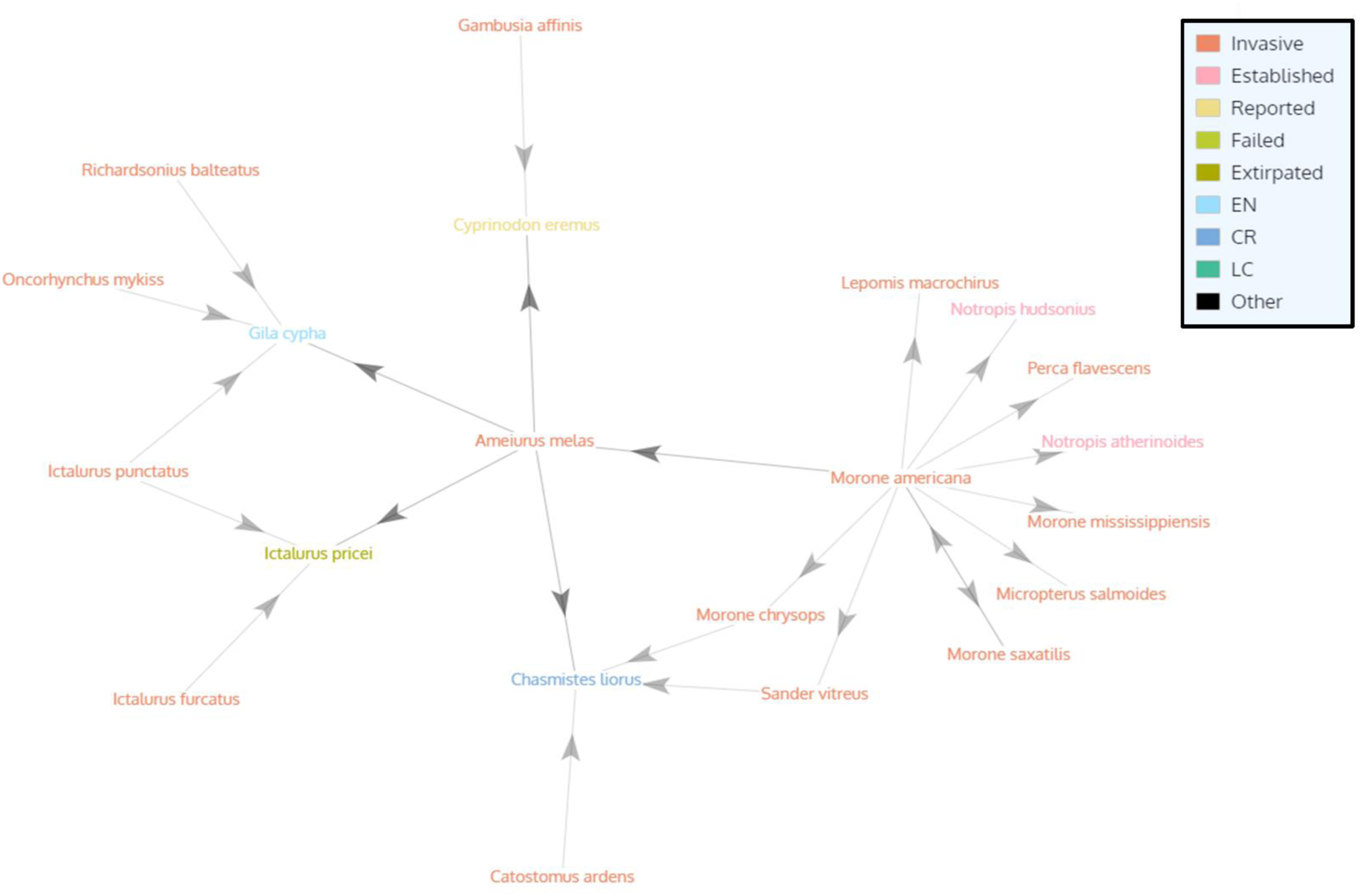
Network showing the direction of impacts of invasive species, as taken from the ITOFF web application. The network is illustrated with *Ameiurus melas* selected as the focal species. The degree of separation between species is set to two, distribution of the species across climate regions is not specified. The web application is illustrated using a colourblind-friendly palette.

### Linking life-history traits and invasive success

Invasive species had greater preferred temperature ranges than impacted species across temperate (13.23 ± 1.07, p = 0.005), subtropical (12.2 ± 1.06, p = 0.001), and tropical (8.38 ± 1.05, p = 0.001) climate regions. Endangered and critically endangered species did not differ significantly from impacted species for preferred temperature range across any of the three climate regions. For all other traits, observed differences between invasive, endangered, critically endangered, and impacted species did not match predictions (Table 1, Figure 2). Further, there was no consistent pattern in trait values between climate zones. The full model outputs are provided in Table A2 (Appendix).

**Figure 2.**
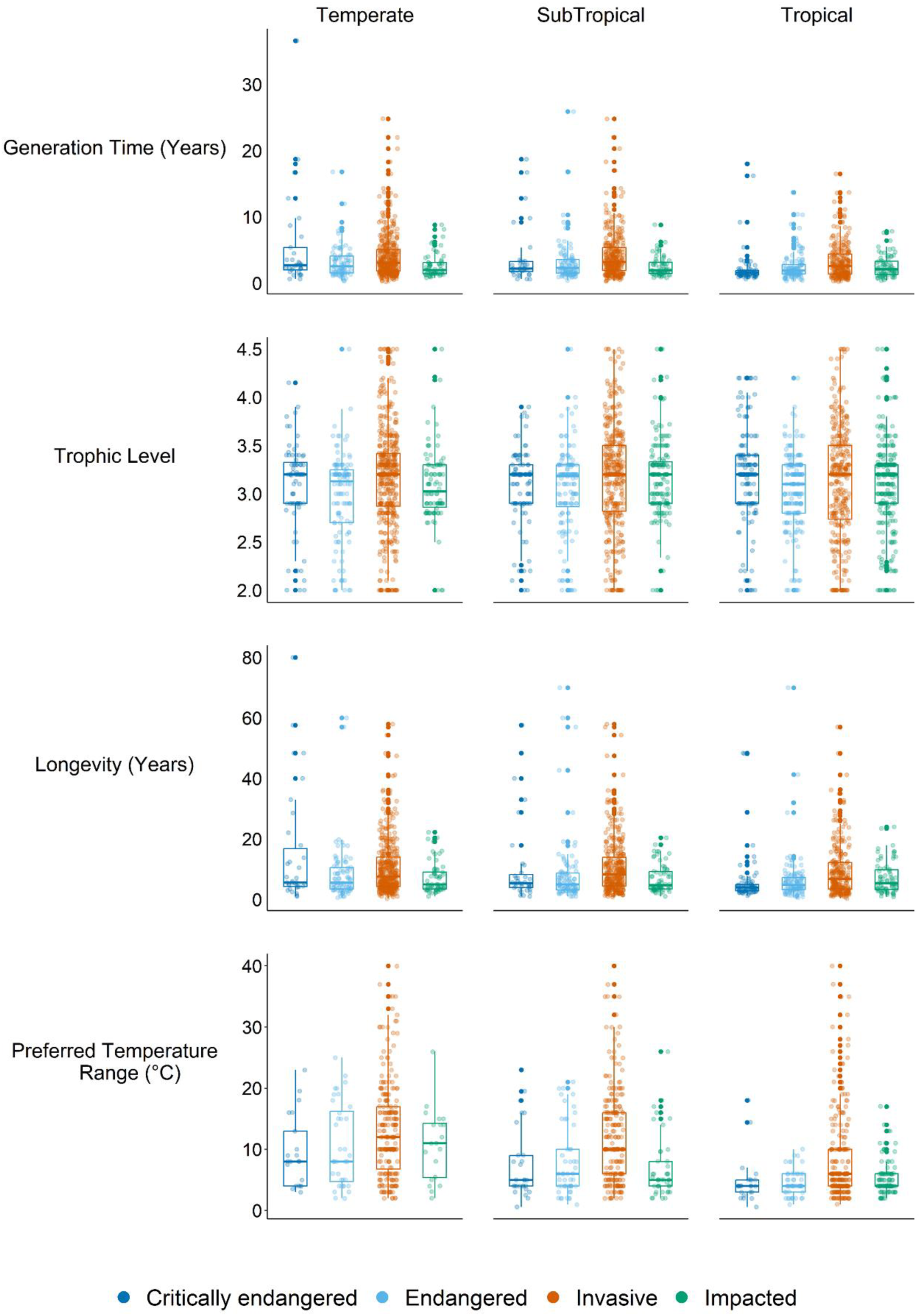
Box and scatter plots showing observed values for generation time (years) (n = 1604), trophic level (n = 2439), longevity (years) (n = 1672), and preferred temperature range (°C) (n = 1074) for invasive species, those species they endanger, critically endanger, and species impacted by invasives but not considered to be at risk of extinction (impacted; control group). Species that occupy multiple climate regions are represented in multiple counts.

**Table 1.** General direction of predicted and observed differences between critically endangered, endangered, and invasive fish species relative to species impacted by invasives but considered least concern in terms of extinction risk (impacted; control group). Green arrows indicate significantly greater trait values for that group, red arrows indicate significantly lower trait values, while a dash indicates no significant differences. Data are split between three major climate regions (Temperate, Subtropical, and Tropical). Asterisks denote statistical significance (*p ≤ 0.05, **p ≤ 0.01, ***p ≤ 0.001). N represents the total number of species included from the corresponding climate region. n indicates the number of species for which data are available for the corresponding trait.

| Climate Region: Temperate<br>(N = 744) |  |  |  |  |  |  |
| --- | --- | --- | --- | --- | --- | --- |
| Trait | Critically endangered |  | Endangered |  | Invasive |  |
|  | Predicted | Observed | Predicted | Observed | Predicted | Observed |
| Generation time<br>(n = 567) | ↑ | ↑ *** | ↑ | — | ↓ | ↑ ** |
| Trophic level<br>(n = 725) | ↓ | — | ↓ | — | — | — |
| Longevity<br>(n = 596) | ↑ | ↑ *** | ↑ | ↑ ** | ↓ | ↑ *** |
| Preferred temperature range<br>(n = 309) | ↓ | — | ↓ | — | ↑ | ↑ ** |

| Climate Region: Subtropical<br>(N = 800) |  |  |  |  |  |  |
| --- | --- | --- | --- | --- | --- | --- |
| Trait | Critically endangered |  | Endangered |  | Invasive |  |
|  | Predicted | Observed | Predicted | Observed | Predicted | Observed |
| Generation time<br>(n = 518) | ↑ | ↑ *** | ↑ | ↑ * | ↓ | ↑ *** |
| Trophic level<br>(n = 779) | ↓ | — | ↓ | ↓ * | — | — |
| Longevity<br>(n = 545) | ↑ | ↑ *** | ↑ | ↑ *** | ↓ | ↑ *** |
| Preferred temperature range<br>(n = 342) | ↓ | — | ↓ | — | ↑ | ↑ *** |

| Climate Region: Tropical<br>(N = 974) |  |  |  |  |  |  |
| --- | --- | --- | --- | --- | --- | --- |
| Trait | Critically endangered |  | Endangered |  | Invasive |  |
|  | Predicted | Observed | Predicted | Observed | Predicted | Observed |
| Generation time<br>(n = 519) | ↑ | — | ↑ | — | ↓ | ↑ *** |
| Trophic level<br>(n = 935) | ↓ | ↑ * | ↓ | — | — | — |
| Longevity<br>(n = 531) | ↑ | ↓ ** | ↑ | — | ↓ | ↑ *** |
| Preferred temperature range<br>(n = 423) | ↓ | — | ↓ | — | ↑ | ↑ *** |

In temperate zones, invasive, endangered, and critically endangered species did not differ from impacted species in terms of mean trophic level (Table 1, Figure 2). Contrary to our predictions, invasive, and critically endangered species had significantly greater generation times than impacted species. Further, invasive, endangered, and critically endangered species all had greater longevity than impacted species (Table 1, Figure 2).

Similar to temperate zones, invasive, endangered, and critically endangered species had greater generation times and longevity than impacted species in subtropical zones (Table 1, Figure 2). Additionally, neither invasive nor critically endangered species differed significantly from impacted species for trophic level (Table 1, Figure 2). Conversely, endangered species had significantly lower trophic level values than impacted species.

In tropical zones, invasive species had significantly greater generation times, longevity, and preferred temperature ranges than impacted species but did not differ significantly for trophic level (Table 1, Figure 2). In contrast, endangered species did not differ significantly from impacted for any of the four traits. Contrary to our predictions, critically endangered species had greater trophic level scores and reduced longevity relative to impacted species (Table 1, Figure 2). Critically endangered species did not differ from impacted species for generation time or temperature range in tropical zones.

### Multiple traits in determining invasive success

Following a divisive cluster analysis based on three traits, some clusters were found to be composed exclusively by invasive species suggesting that combined traits may be linked to invasive success (Figure 3A, green circles). However, other clusters contained invasive, impacted and/or endangered species (Figure 3A, yellow circles). Accordingly, multiple invasive species have greater similarity to impacted and endangered species as opposed to other invasives. Following phylogenetic analyses of the same species, we identified genera that contain a mix of invasive, impacted and/or endangered species (Figure 3B, blue circles). However, a genera (*Pterygoplichthys*) containing only invasive species was also identified (Figure 3B, red circle).

**Figure 3.**
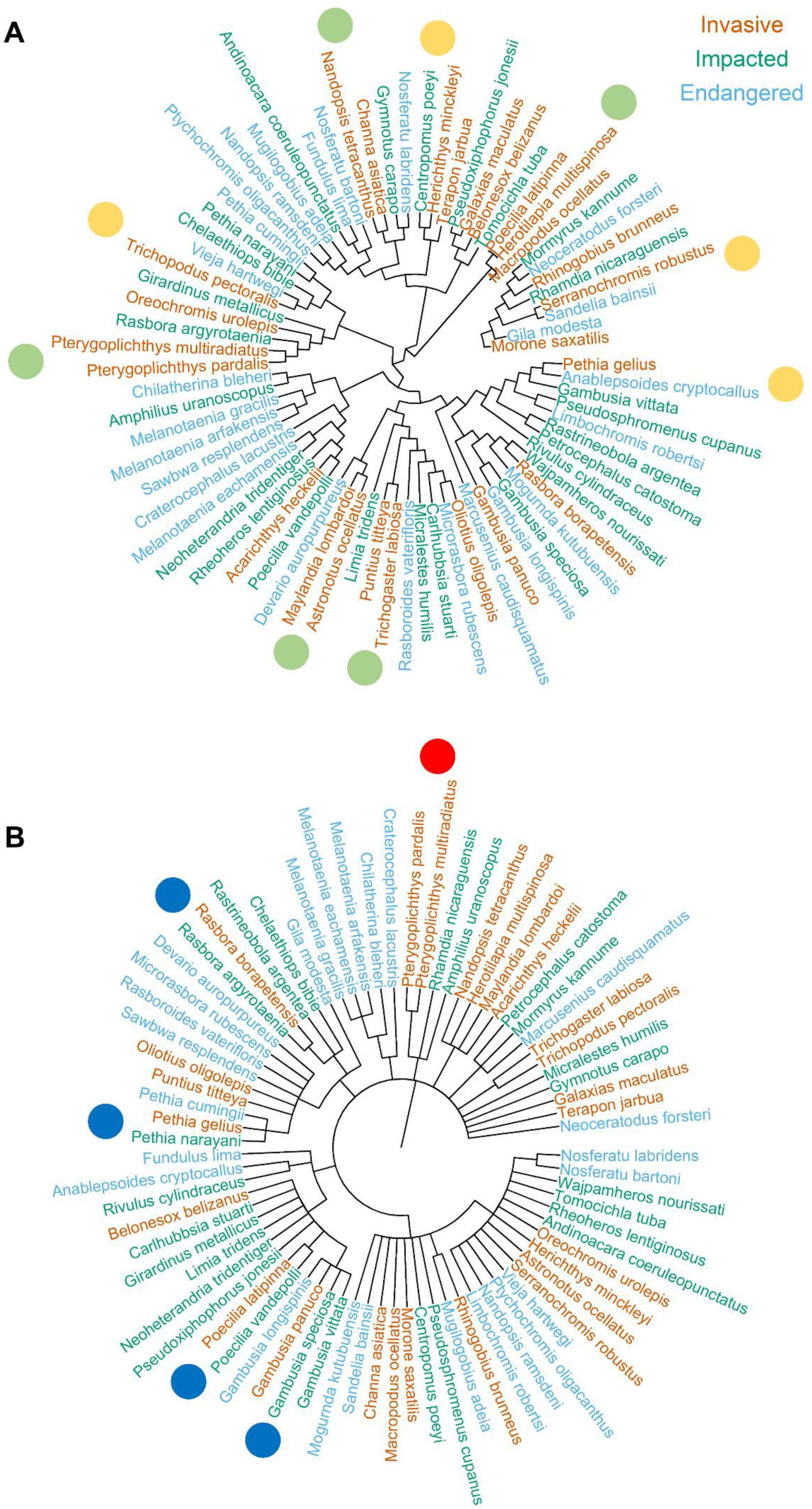
(A) Hierarchical cluster (using the Divisive ANAlysis [DIANA] clustering algorithm) based on trait values for age of sexual maturity, trophic level, and temperature range for a random subset of 75 tropical species in the ITOFF database (25 invasive species, 25 species endangered due to the spread of invasive species, and 25 species impacted by invasives but not endangered [impacted]). Green circles indicate clusters composed of exclusively invasive species. Yellow circles indicate clusters containing invasive, impacted and/or endangered species. (B) Taxonomic tree constructed for the same species pool used in (A). Red circles indicate genera containing only invasive species. Blue circles indicate genera containing invasive, impacted and/or endangered species.

## DISCUSSION

ITOFF offers a unique and accessible platform to comprehensively address key questions in invasion biology. ITOFF data provides strong evidence that classical predictions for life-history traits facilitating invasive success do not hold for freshwater fish on the major climate zone scale (Capellini *et al*., 2015; Chapple, Simmonds, and Wong, 2012; Crowder and Snyder, 2010). While invasive freshwater fish species are characterized by greater preferred temperature ranges, they cannot be reliably identified by generation time, trophic level, longevity, or multiple trait analyses. It is therefore essential that we shift the focus of invasive species management towards a holistic approach that considers the species (Chapple, Simmonds, and Wong, 2012), environment (Pyšek *et al*., 2020), and evolutionary history (Mazzamuto *et al*., 2016).

### ITOFF Database

The ITOFF database reveals key gaps in data availability amongst invasive species and those species they impact. For example, there is data inequality amongst non-native species. Specifically, trait data are unavailable for many failed invaders. Nonetheless, comparisons between non-native groups (i.e. invasive, and failed invaders) will allow the identification of traits that facilitate invasive success during one or more stages of the invasion process (Capellini *et al*., 2015). Future research should target both successful and unsuccessful invasive freshwater fish species. Additionally, by using ITOFF it was possible to identify key data gaps concerning the distribution of invasive species across climate regions. One of the major factors affecting the spread of non-native species is tropicalization (Osland *et al*., 2021). High latitude regions are therefore hot spots for the spread of non-native species (Goldsmit *et al*., 2020; Hughes *et al*., 2020). However, data on freshwater invasions in these regions are limited relative to tropical, subtropical and temperate regions. It is therefore critical that invasive species research is distributed evenly across climate zones.

Another unique addition provided by ITOFF to invasion biology research is the visualization of impacts associated with invasive species. This is an accessible tool with applications for academic research, citizen science, and education. Networks highlight the complexity of impacts associated with invasive species and can be used to guide invasive species management and identify target study species.

### Linking life-history traits and invasive success

Collectively, the results of our trait analyses provide weak evidence to support the classical predictions that life-history traits promote invasive success (Capellini *et al*., 2015; Chapple, Simmonds, and Wong, 2012; Crowder and Snyder, 2010). The influence of environmental variability and propagule pressure on invasive success may explain the failure to detect life-history traits that ubiquitously promote invasive success (Moyle and Light, 1996; Tabak, Webb, and Miller, 2018). However, a key finding of this study is the greater preferred temperature ranges for invasive species across temperate, tropical, and subtropical climate zones. This result confirms predictions that broader abiotic tolerances increase the likelihood of invasive success by facilitating survival in a greater range of novel environments (Zerebecki and Sorte, 2011; Bates *et al*., 2013). Nonetheless, it should be noted that there is large overlap in trait values amongst all treatment groups for all traits, including preferred temperature range (Figure 2).

### Multiple traits in determining invasive success

The confounding results of the hierarchical cluster and phylogenetic analyses stress the complexity of the invasion process and suggest that both life-history traits and context are key in determining invasive success. These results prompt discussion into the role of ancestral traits in determining invasive success and highlight the need for targeted research of closely related species that differ in invasive status.

Overlap of trait values between invasive species and those species they impact limits the successful identification of potential invasive species on the climatic zone or global scale. This key result is in line with previous conceptual research which predicted that ubiquitous invasive traits are unlikely to be identified (Moyle and Light, 1996). Universal invasive species management is likely to be less efficient than localized management strategies and we advise that this be reflected in policy. Successful prediction of invasion on reduced spatial scales is possible with detailed knowledge of the biotic and abiotic conditions of an ecosystem (Kolar and Lodge, 2002; Marchetti, Moyle and Levine, 2004; Vila-gispert, Alcaraz and Garcia-Berthou, 2005). However, local scale approaches to invasive species management are work intensive. Nonetheless, as current ability to forecast aquatic species invasions on the large spatial scale remains limited, localised management remains the most viable approach (Moyle and Light, 1996).

### Future directions

The accurate prediction of future invasive species is a central component of governmental non-native species management. In the United Kingdom, ‘The Great Britain Invasive Non-native Species Strategy’ (Department for Environment Food and Rural Affairs [DEFRA], 2015) is used as a strategic framework for current management (e.g. ‘River basin management plans’; Environment Agency, 2022). Horizon scanning is the only predictive management measure named in the DEFRA framework, and is largely reliant on the use of ‘biological and ecological criteria’ to identify species with a high-risk of becoming invasive in the UK. The results of this study demonstrate that predictive frameworks must consider ecosystem characteristics, evolutionary history and contemporary evolution in combination with life-history traits for improved accuracy of invasive species forecasts. We argue that urgent research concerning the relationship of life-history traits, evolutionary history, rapid adaptation and invasive success is needed. ITOFF opens the opportunity to test the above avenues of research, which ultimately will further our understanding of the factors contributing to successful invasion in freshwater fish, a model taxon for the study of invasion biology.

## Supporting information

Variables Description

## Data Availability Statement

The ITOFF core dataset is available at https://doi.org/10.5281/zenodo.10135093

## Acknowledgements

The authors wish to acknowledge the U.S. Geological Survey, the Invasive Species Specialist Group, the IUCN Red List of Threatened Species, FishBase, and CABI for the provision of data that have made the ITOFF project possible. M.B. is supported by a FCT/MCTES (Fundação para a Ciência e Tecnologia/Ministério da Ciência, Tecnologia e Ensino Superior) postdoctoral fellowship (<u>SFRH/BPD/82259/2011</u>). M.B. also acknowledges the financial support provided by CESAM (<u>UIDP/50017/2020+UIDB/50017/2020</u>).

## APPENDIX

### ITOFF usage notes

The core ITOFF dataset can be downloaded from the Zenodo public repository (https://doi.org/10.5281/zenodo.10135093) and should be referenced by citing the present paper. Exemplary R code and a document describing the variables included in the core ITOFF dataset are also available via the Zenodo public repository. Additionally, a dynamic, continuously updated, version of the ITOFF database can be downloaded from the ITOFF project website (https://itoff-dataset.wp.st-andrews.ac.uk). Data are organised such that the dataset is easy to use and serves as a readily available tool for invasion biology research. The web application can be accessed via the ITOFF project website (https://itoff-dataset.wp.st-andrews.ac.uk/webapplication/). Guidance on how to use the web application is also provided on the project website.

The primary intended use of the dataset is the identification of shared invasive traits amongst freshwater fish. Traits can be identified through broad-scale comparisons made between invasive, critically endangered, endangered, and impacted species. Once identified, these traits will improve current ability to forecast species invasions and highlight ‘high-risk’ species with great invasive potential. The same data offer insights into traits that may increase a native species’ susceptibility to invasion and can therefore be used to inform policy, conservation, and invasive species management. Future studies can address differences in invasive traits between climate regions. This research is of great importance given the current knowledge gap concerning how climate change will impact the outcome of future species invasions. Additionally, using the provided taxonomic information, future studies can untangle the role of adaptation, chance, and evolutionary history in the invasion process (Figure 1A, Appendix). Further, cluster analyses can investigate similarity across multiple traits for invasive, endangered, and impacted species (Figure 1B, Appendix). Cluster analyses will be important to invasive trait identification as invasive propensity is not always tied to individual traits but arises from a combination of multiple traits (Grabowski, Bacela and Konopacka, 2007).

**Table A1.** Descriptions of the life-history traits included in the ITOFF database and evidence for their link to invasive success.

| Trait | Description | Link to invasive success | References |
| --- | --- | --- | --- |
| **Reproductive guild (included as mode of reproduction, mode of fertilization, and extent of parental care)** | Species are categorised as non-guarders, guardians, and bearers. Bearers include livebearers, mouthbrooders, or any species that utilises a brood pouch. Fish that actively defend externally laid eggs are guardians. Non-guarders include egg scatterers and any species for which neither parent actively defends eggs. Additional information is provided in a supplementary column (i.e. mouth | Greater parental care promotes colonisation success, especially in harsh environments. Further, establishment success was correlated with greater parental investment in Californian fishes and a global study using 14 fish species. Further, live birth is linked to increased offspring survival, colonisation success, and is advantageous when mating opportunities are limited (i.e. when | (Wourms and Lombardi, 1992; Marchetti, Moyle and Levine, 2004; Drake, 2007; Deacon, Ramnarine, and Magurran, 2011; Wong, Meunier and Kölliker, 2013) |
|  | brooding, nesting, internal live-bearing). | population sizes are limited following introduction). |  |
| **Absolute fecundity** | Total number of eggs in an average female (for egg-layers), or the average, total number of offspring produced over the entire lifetime of livebearers. | Greater reproductive output is positively associated with mammalian invasion success on the global scale | (Capellini *et al.*, 2015) |
| **Brood size** | The number of offspring produced, or the number of eggs laid at one discrete time | Production of larger broods reduces the risk of extinction for small populations and is linked to non-native mammalian establishment success | (Pimm, 1991; Capellini *et al.*, 2015) |
| **Age of sexual maturity** | The average age (years) at which an individual is able to successfully reproduce. Provided as a mean value across sex in the ITOFF database | Early maturity is linked to rapid population growth, establishment success, and improved resistance to demographic stochasticity and anthropogenic management strategies in the invasive common carp, *Cyprinus carpio*. Conversely, non-native fish species of Catalan streams are characterised by late maturity relative to native species | (Vila-gispert, Alcaraz and Garcia-Berthou, 2005; Winker *et al.*, 2011) |
| **Generation time** | The average age (years) of parents at the time their first offspring are born | Short generation times promote rapid population growth, colonisation success, and facilitates rapid adaptation | (Rosenthal *et al.*, 2021) |
| **Trophic level and feeding type** | Trophic level is a value estimated by FishBase to show the 'rank of a species in a food web'. Defined as 1 + the mean trophic level of the focal species' food items (Froese and Pauly, 2022). Under feeding type, fish are categorised by mode of feeding (e.g. grazer or predator). | Generalist diets allow for behavioural flexibility in novel environments and increase the competitive ability of non-native species | (Crowder and Snyder, 2010) |
| **Relative brain weight** | Given as brain weight (mg) divided by body weight (g). | Larger brains may facilitate greater behavioural flexibility and promote survival in novel environments. Brain size is correlated with invasive success in birds, mammals, amphibians, and reptiles | (Sol *et al.*, 2008; Vall-Llosera and Sol, 2009; Amiel, Tingley and Shine, 2011) |
| **Longevity (included as maximum age in years, maximum weight, and maximum length)** | The maximum age, weight, or length that an individual of each species can reach. Reported as the mean value of the approximate age, weight, or length of the oldest known individuals amongst populations. | Though life-history strategies are generally acknowledged to be an important factor in determining invasive success, there is debate as to whether or not invasive species are more often characterised by fast or slow life-histories. Fast life-histories are predicted to promote rapid population growth and establishment success. However, slow life-histories can also increase establishment success in a wider range of environmental conditions by prioritising survival and delaying reproduction | (Sol *et al.*, 2012) |
| **Temperature range (included as both preferred and tolerable)** | Preferred temperature range: the range of temperatures over which the species can survive indefinitely. Reported in ITOFF as the range of water temperatures experienced throughout the species range.<br>Tolerable temperature range: the range of temperatures over which individuals of the species can survive brief exposure | Wider abiotic tolerances increase the likelihood of survival within a wider range of novel environments | (Zerebecki and Sorte, 2011; Bates *et al.*, 2013) |

**Table A2.**
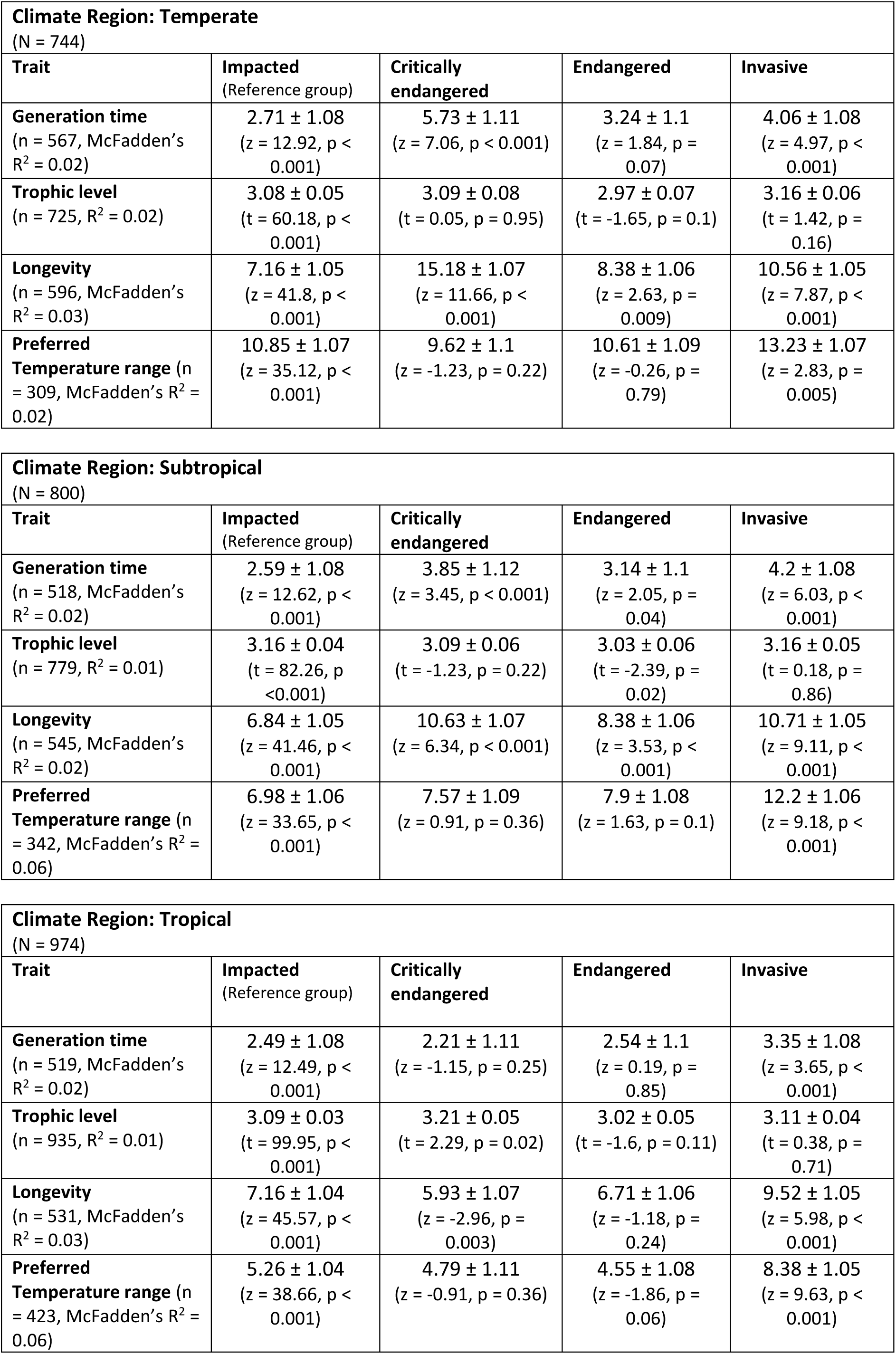
Summary of the results for generalized linear models investigating differences in life-history trait values between invasive species, species impacted by invasive species but not facing an elevated risk of extinction (impacted), endangered species that are threatened by invasive species, and critically endangered species that are threatened by invasive species. Results are shown for four life-history traits shown to be critical to invasive success. Data are split between major climate regions. Species present in multiple climate regions are represented in multiple counts. N represents the total number of species included for the corresponding climate region. n indicates the number of species for which data are available for the corresponding trait. Overall model adjusted R^2^ is reported for models with Gaussian error distributions, McFadden’s pseudo-R^2^ is reported for models with Poisson error distributions.

| Climate Region: Temperate<br>(N = 744) |  |  |  |  |
| --- | --- | --- | --- | --- |
| Trait | Impacted<br>(Reference group) | Critically<br>endangered | Endangered | Invasive |
| **Generation time**<br>(n = 567, McFadden's<br>R<sup>2</sup> = 0.02) | 2.71 ± 1.08<br>(z = 12.92, p <<br>0.001) | 5.73 ± 1.11<br>(z = 7.06, p < 0.001) | 3.24 ± 1.1<br>(z = 1.84, p =<br>0.07) | 4.06 ± 1.08<br>(z = 4.97, p <<br>0.001) |
| **Trophic level**<br>(n = 725, R<sup>2</sup> = 0.02) | 3.08 ± 0.05<br>(t = 60.18, p <<br>0.001) | 3.09 ± 0.08<br>(t = 0.05, p = 0.95) | 2.97 ± 0.07<br>(t = -1.65, p = 0.1) | 3.16 ± 0.06<br>(t = 1.42, p =<br>0.16) |
| **Longevity**<br>(n = 596, McFadden's<br>R<sup>2</sup> = 0.03) | 7.16 ± 1.05<br>(z = 41.8, p <<br>0.001) | 15.18 ± 1.07<br>(z = 11.66, p <<br>0.001) | 8.38 ± 1.06<br>(z = 2.63, p =<br>0.009) | 10.56 ± 1.05<br>(z = 7.87, p <<br>0.001) |
| **Preferred**<br>**Temperature range** (n<br>= 309, McFadden's R<sup>2</sup> =<br>0.02) | 10.85 ± 1.07<br>(z = 35.12, p <<br>0.001) | 9.62 ± 1.1<br>(z = -1.23, p = 0.22) | 10.61 ± 1.09<br>(z = -0.26, p =<br>0.79) | 13.23 ± 1.07<br>(z = 2.83, p =<br>0.005) |

| Climate Region: Subtropical<br>(N = 800) |  |  |  |  |
| --- | --- | --- | --- | --- |
| Trait | Impacted<br>(Reference group) | Critically<br>endangered | Endangered | Invasive |
| **Generation time**<br>(n = 518, McFadden's<br>R<sup>2</sup> = 0.02) | 2.59 ± 1.08<br>(z = 12.62, p <<br>0.001) | 3.85 ± 1.12<br>(z = 3.45, p < 0.001) | 3.14 ± 1.1<br>(z = 2.05, p =<br>0.04) | 4.2 ± 1.08<br>(z = 6.03, p <<br>0.001) |
| **Trophic level**<br>(n = 779, R<sup>2</sup> = 0.01) | 3.16 ± 0.04<br>(t = 82.26, p<br><0.001) | 3.09 ± 0.06<br>(t = -1.23, p = 0.22) | 3.03 ± 0.06<br>(t = -2.39, p =<br>0.02) | 3.16 ± 0.05<br>(t = 0.18, p =<br>0.86) |
| **Longevity**<br>(n = 545, McFadden's<br>R<sup>2</sup> = 0.02) | 6.84 ± 1.05<br>(z = 41.46, p <<br>0.001) | 10.63 ± 1.07<br>(z = 6.34, p < 0.001) | 8.38 ± 1.06<br>(z = 3.53, p <<br>0.001) | 10.71 ± 1.05<br>(z = 9.11, p <<br>0.001) |
| **Preferred**<br>**Temperature range** (n<br>= 342, McFadden's R<sup>2</sup> =<br>0.06) | 6.98 ± 1.06<br>(z = 33.65, p <<br>0.001) | 7.57 ± 1.09<br>(z = 0.91, p = 0.36) | 7.9 ± 1.08<br>(z = 1.63, p = 0.1) | 12.2 ± 1.06<br>(z = 9.18, p <<br>0.001) |

| Climate Region: Tropical<br>(N = 974) |  |  |  |  |
| --- | --- | --- | --- | --- |
| Trait | Impacted<br>(Reference group) | Critically<br>endangered | Endangered | Invasive |
| **Generation time**<br>(n = 519, McFadden's<br>R<sup>2</sup> = 0.02) | 2.49 ± 1.08<br>(z = 12.49, p <<br>0.001) | 2.21 ± 1.11<br>(z = -1.15, p = 0.25) | 2.54 ± 1.1<br>(z = 0.19, p =<br>0.85) | 3.35 ± 1.08<br>(z = 3.65, p <<br>0.001) |
| **Trophic level**<br>(n = 935, R<sup>2</sup> = 0.01) | 3.09 ± 0.03<br>(t = 99.95, p <<br>0.001) | 3.21 ± 0.05<br>(t = 2.29, p = 0.02) | 3.02 ± 0.05<br>(t = -1.6, p = 0.11) | 3.11 ± 0.04<br>(t = 0.38, p =<br>0.71) |
| **Longevity**<br>(n = 531, McFadden's<br>R<sup>2</sup> = 0.03) | 7.16 ± 1.04<br>(z = 45.57, p <<br>0.001) | 5.93 ± 1.07<br>(z = -2.96, p =<br>0.003) | 6.71 ± 1.06<br>(z = -1.18, p =<br>0.24) | 9.52 ± 1.05<br>(z = 5.98, p <<br>0.001) |
| **Preferred**<br>**Temperature range** (n<br>= 423, McFadden's R<sup>2</sup> =<br>0.06) | 5.26 ± 1.04<br>(z = 38.66, p <<br>0.001) | 4.79 ± 1.11<br>(z = -0.91, p = 0.36) | 4.55 ± 1.08<br>(z = -1.86, p =<br>0.06) | 8.38 ± 1.05<br>(z = 9.63, p <<br>0.001) |

#### ITOFF Supplementary R code

R code for basic analysis of the ITOFF database. This script will be provided in unison with the ITOFF database following publication of the ITOFF data paper.

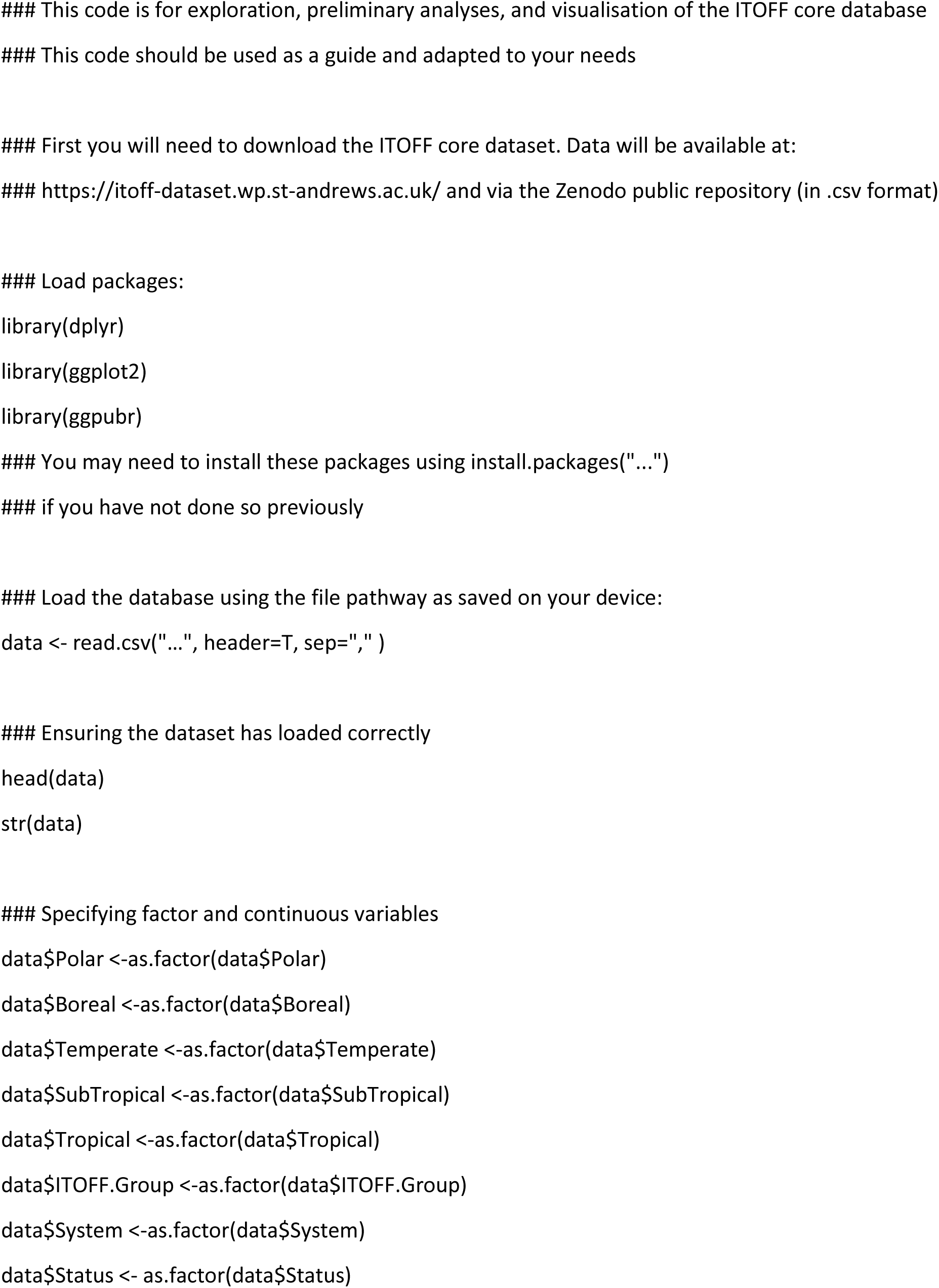
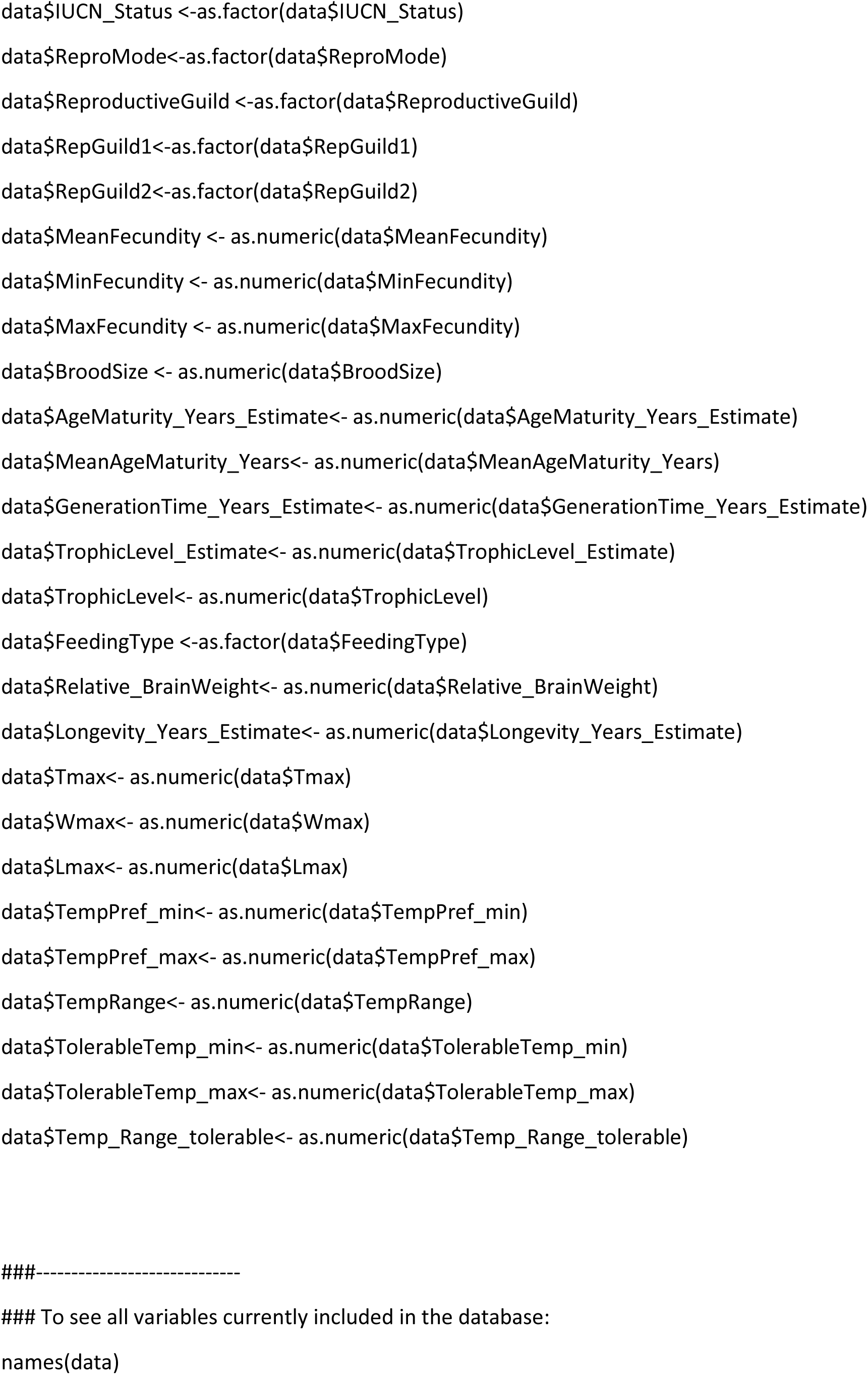
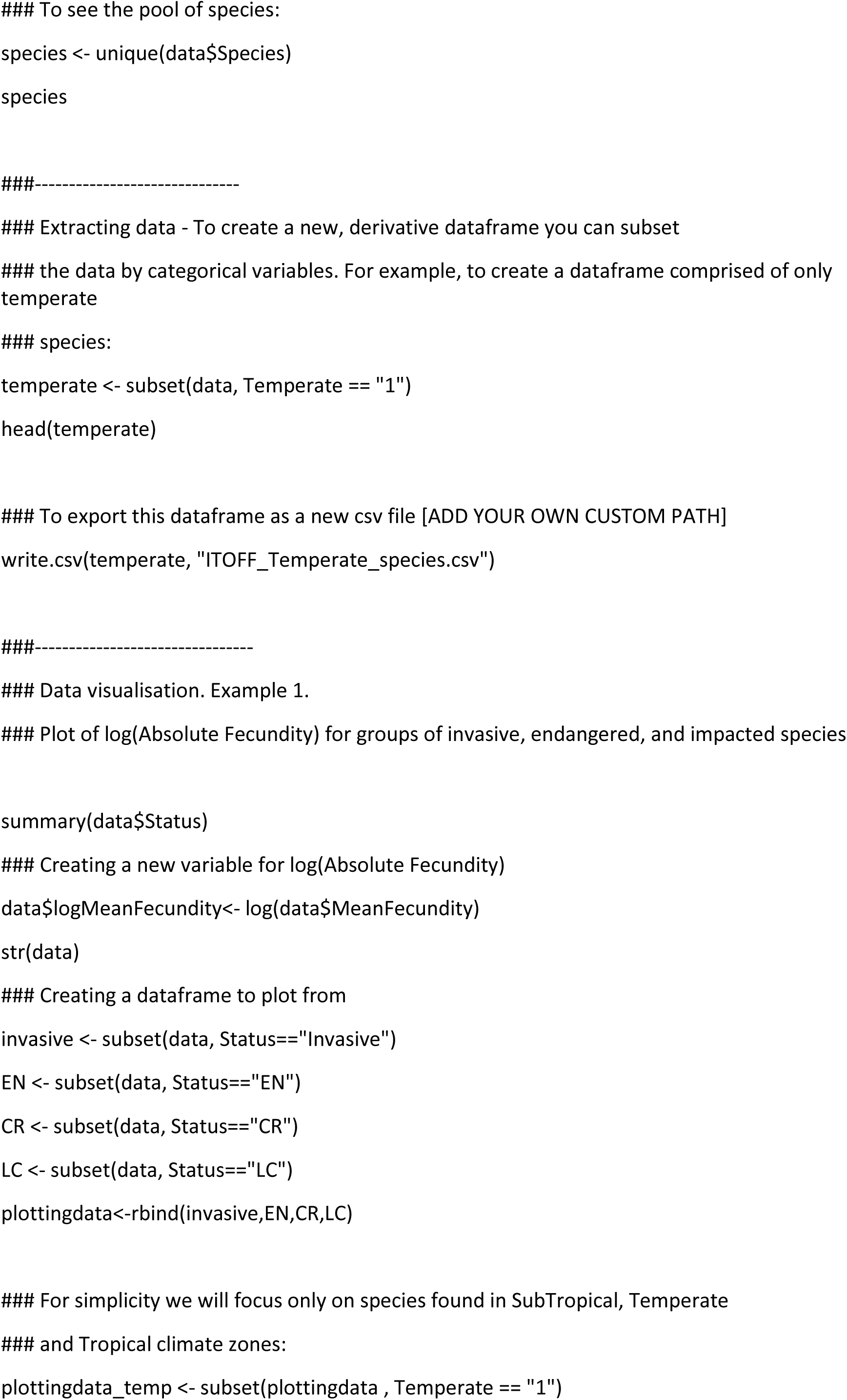
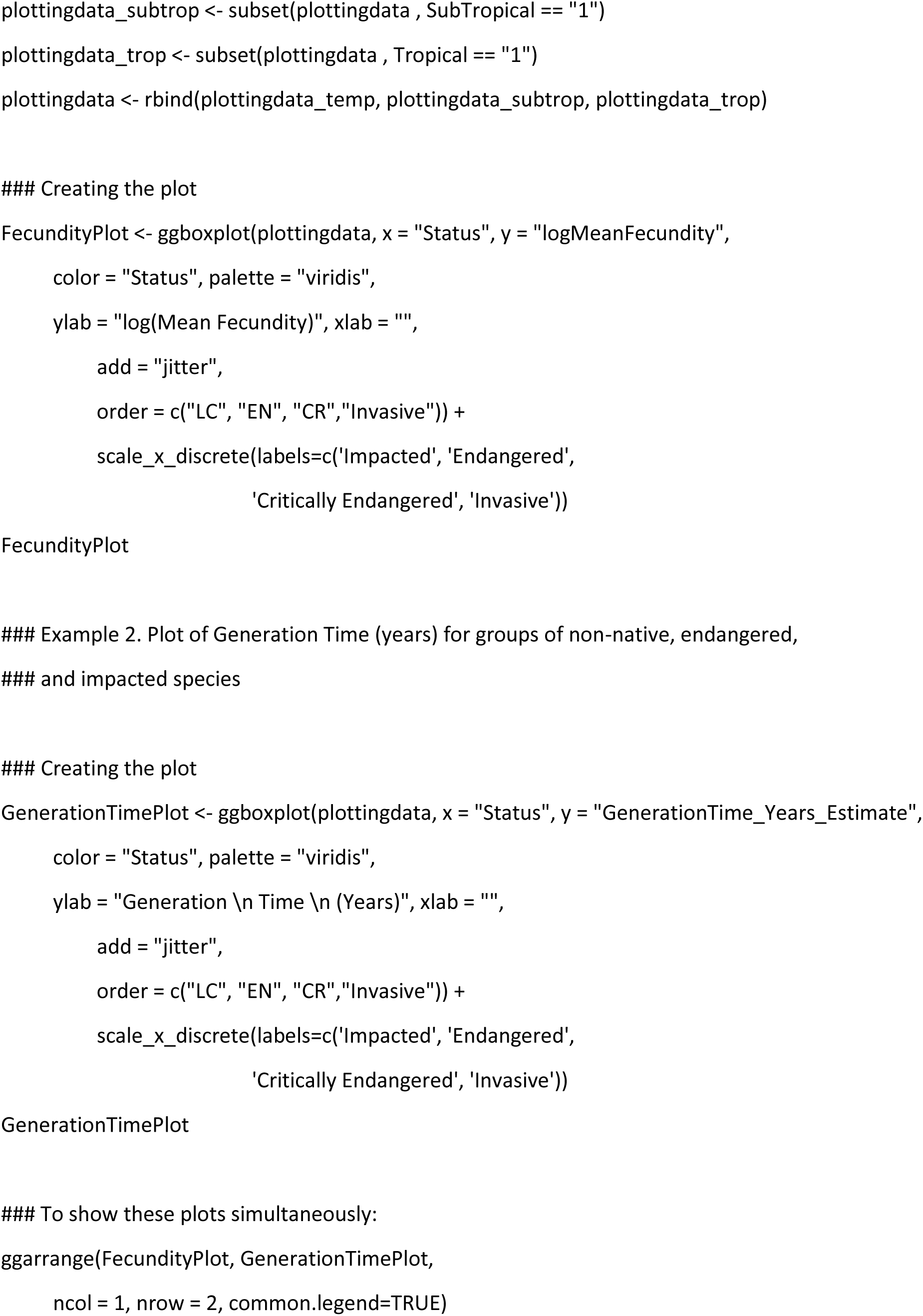
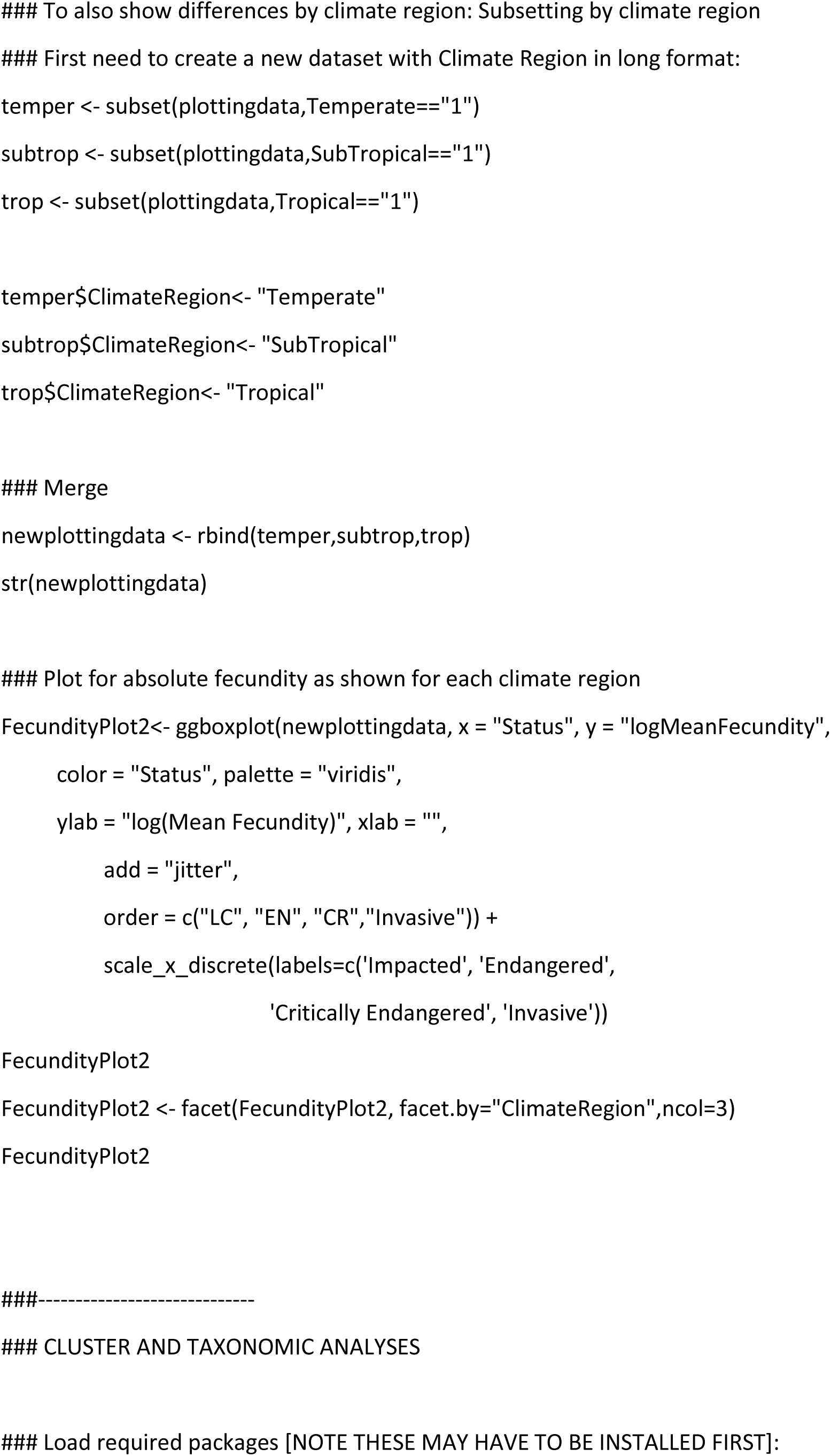
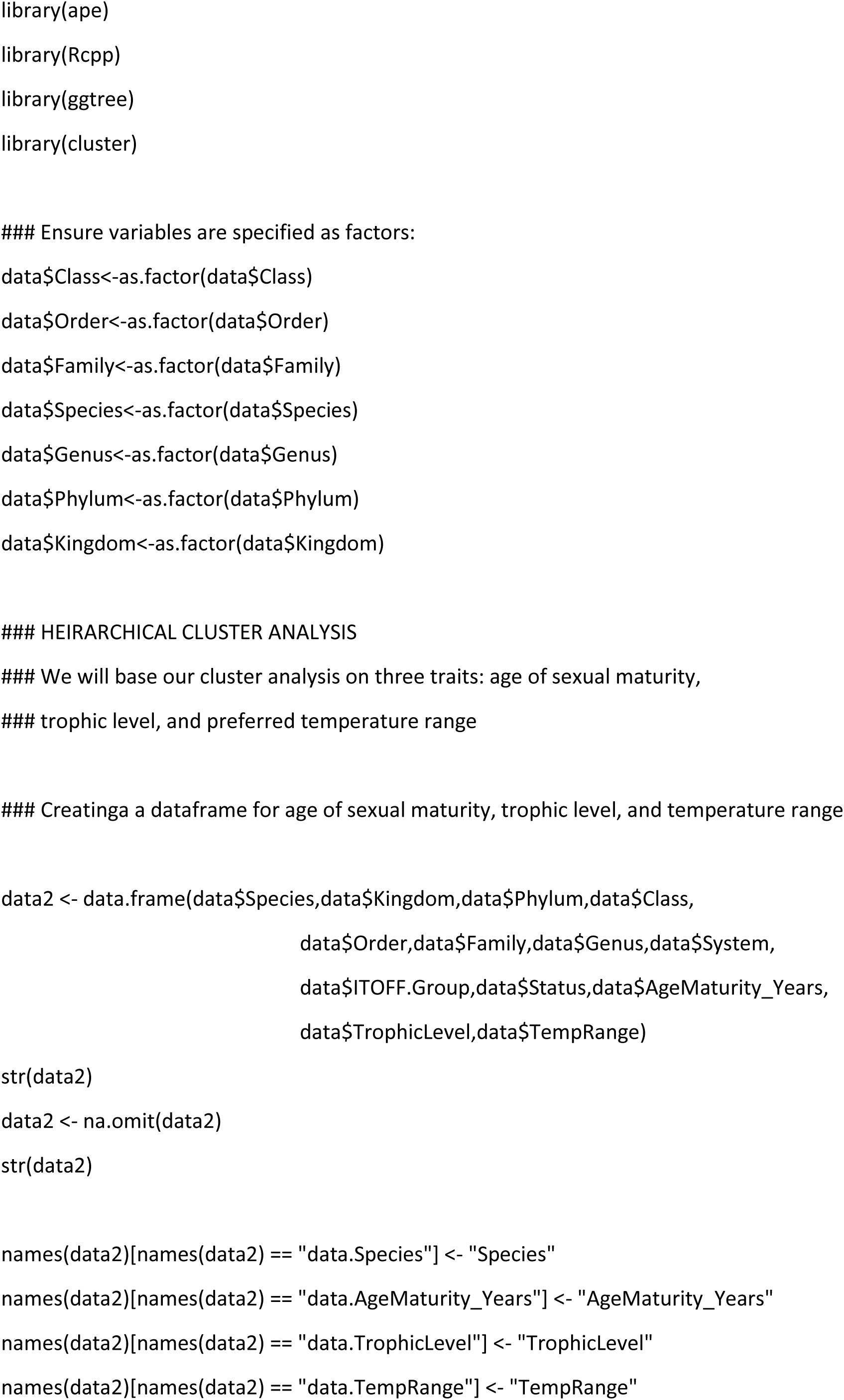
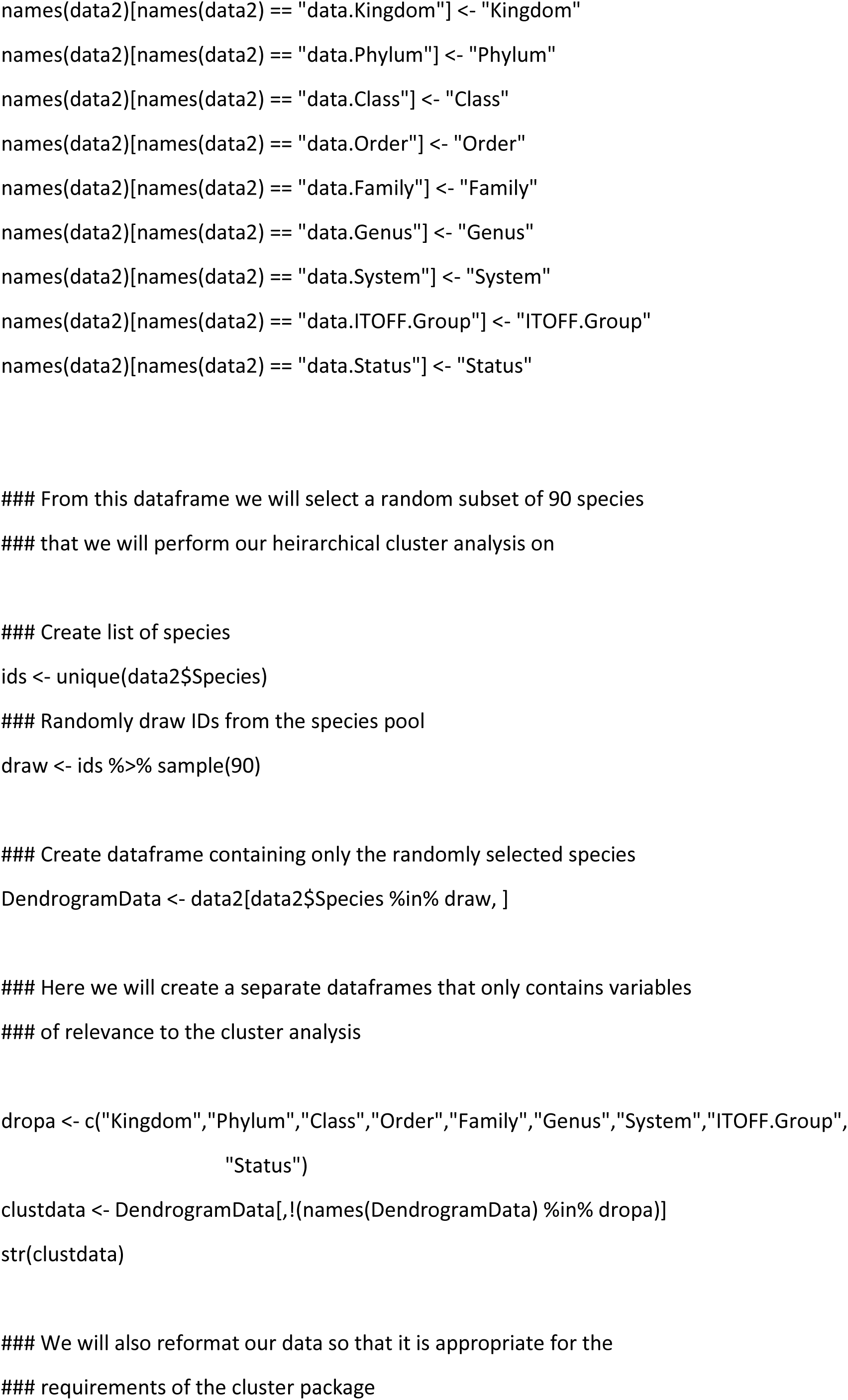
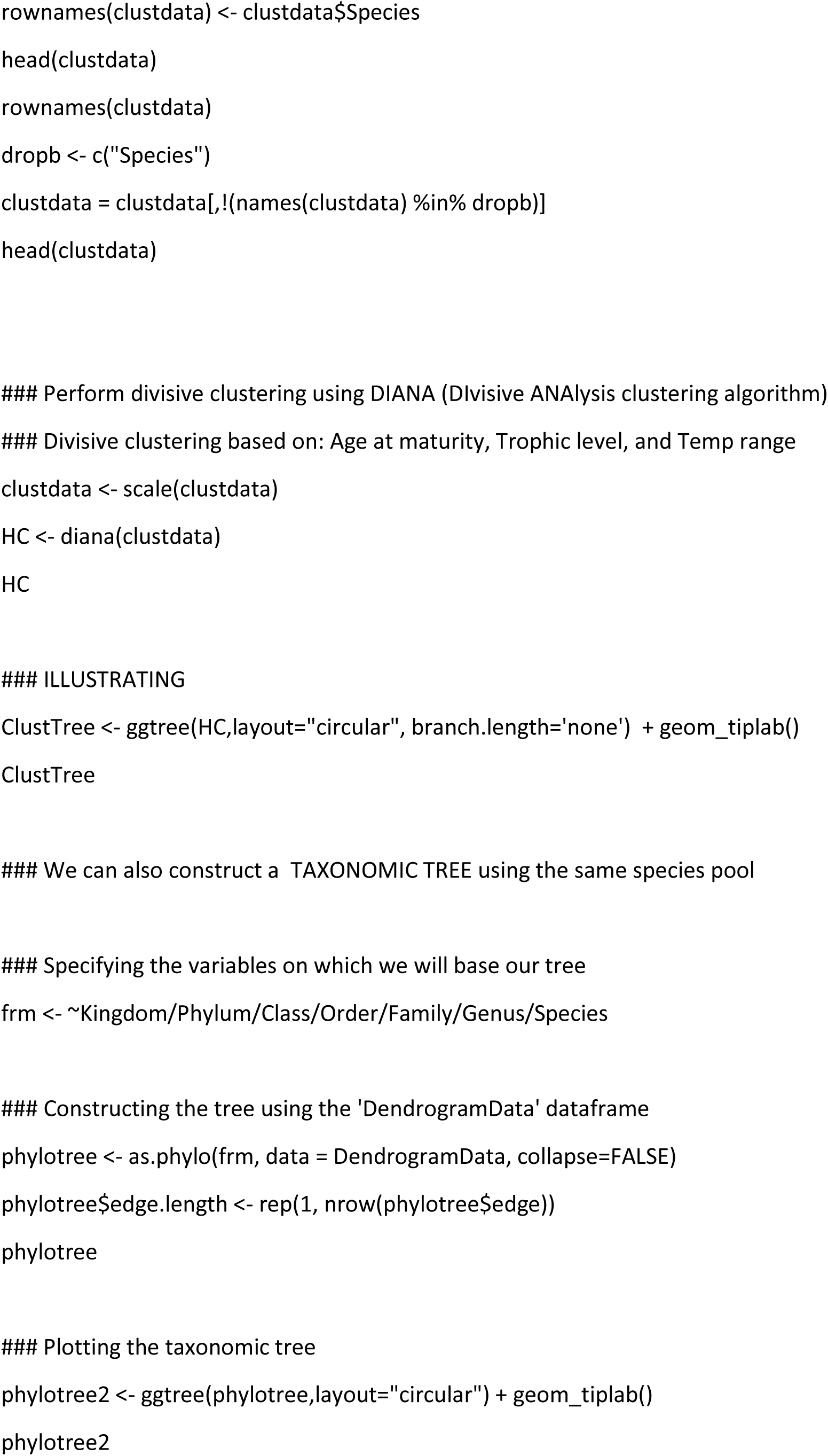
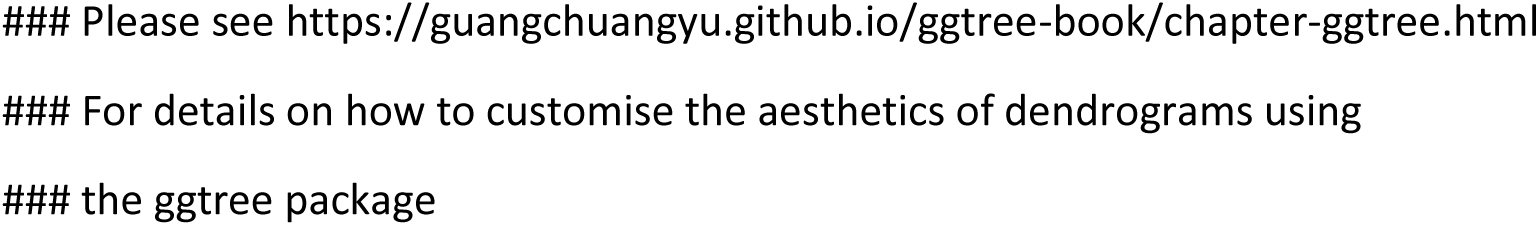

