## Supplementary material for "Invasive traits of freshwater fish database (ITOFF)": Variables Description

***DESCRIPTION OF ITOFF DATA***

***Species:*** Taxonomic binomial nomenclature

***SpecCode:*** The unique species code as specified by FishBase

***CommonName:*** The colloquial name of the species

***ITOFF.Group:*** 1- Non-native, 2 – Endangered (with invasive species considered a threat), 3 – Critically Endangered (with invasive species considered a threat), 4 – Impacted by invasive species but not considered to be at risk of extinction, 5 – Locally threatened by invasive species, 6 – Impacted by invasive species and IUCN Red List Status is unknown, vulnerable, near threatened, or extinct.

***Kingdom, Phylum, Class, Order, Family, Genus:*** Taxonomic rank as in accordance with FishBase as of 01/10/2023

***System:*** Ecosystem(s) the species can be found in, separated by underscore e.g. Freshwater_Brackish, Freshwater_Brackish_Marine.

***Group:*** Recorded as ‘Native’ for speices with no reported introduced populations, and ‘Nonnative’ for species with known introductions regardless of the introduction outcome.

***Status:*** For native species, Status is the same as their IUCN Red List of Threatened Species Status. For invasive species status is one of: ‘Reported’, ‘Established’, ‘Invasive’, ‘Extirpated’, and ‘Failed’. Reported is the default for all non-native species. Established refers to a reported species that has successfully reproduced within its new range. Invasive refers to established species that have or have the potential for any negative (environmental or economic) impact. Extirpated refers to species that initially survived within their new range but were then eliminated by human action. Failed refers to species introduced to non-native ecosystems that do not successfully reproduce.

***EstabWildSucces:*** The proportion of introduced populations that have survived to establishment (instances where survival is unknown or assumed have been omitted). This is based only on the introductions recorded in FishBase and are not representative of all introductions for the species – therefore the species may have a score of 0 here but be a successful invasive elsewhere (this simply indicates that the invasive population has not been reported in FishBase).

***Means_Introduction:*** Modes of introduction to non-native ecosystems. Can be multiple (e.g. “Aquaria, Fish farms”). ‘Passive’ refers to species that dispersed into new ecosystems following the connection of two bodies of freshwater (lakes, streams, rivers etc.) e.g. through canals. Fishing recreation’ refers to species intentionally stocked as gamefish or for any fishing recreation - this also includes species introduced as “forage-fish” which are species introduced for the purposes of feeding the more desirable (often piscivorous) gamefish. ‘Unintentional recreation’ is if the species was introduced unintentionally e.g. it was being used as bait by a fisherman in a river it does not naturally occur in and escaped.

***IUCN_Status:*** Status of the species as reported by the IUCN Red List of Threatened Species.

***Polar, Boreal, Temperate, SubTropical, Tropical:*** Recorded as 1 if the species can be found in the respective climate zone and 0 if the species has not been recorded within the respective climate zone.

***Impacting:*** If the species is known to impact another the impacted species name is given and the type of impact recorded in brackets, multiple impacts on different species are separated by ; (e..g. Archoplites interruptus (Competition); Acheilognathus longipinnis (Predation)). Only species-level impacts are recorded i.e. impacts reported for a genera or any other taxonomic rank will not be shown here.

***Impacted_by:*** As for ‘Impacting’ but reversed.

***ReproMode:*** Species are categorized as dioecious, parthenogenetic, or as having a protandy/protogyny mating system.

***Fertilization:*** Species are characterized by mode of fertilization e.g. internal, external, in-mouth, or via a brood pouch or other structure.

***MatingSystem:*** Categorized as monogamy, polyandry, or polygyny.

***RepGuild1:*** Species are broadly categorized as bearers, guarders, non-guarders,

***RepGuild2:*** More detail on the species reproductive guild, e.g. live-bearer, external brooders, nesters, or scatterers.

***Spawning:*** Character variable describing the frequency of spawning over a year.

***Mean-, min-, max-Fecundity:*** The total number of eggs or live-offspring produced by 1 female of the species over the course of its entire lifetime.

***BroodSize:*** The number of eggs laid, or offspring birthed in one spatially distinct event.

***MeanAgeMaturity_Years:*** Average age (in years) at which individuals of the focal species reach sexual maturity (recorded as the average across males and females). As based on observations.

***AgeMaturity_Years_Estimate:*** As above but as estimated by the FishBase life-history tool. Estimated values are not provided if age of maturity has been observed.

***GenerationTime_Years_Estimate:*** An estimate of the species’ generation time as produced by the FishBase life-history tool.

***TrophicLevel:*** Unique score of the species’ trophic level as given by FishBase.

***TrophicLevel_Estimate:*** As above but estimated for the species using closest relatives.

***FeedingType:*** Categorization of mode of feeding, e.g. predator, plankton feeding, variable, or grazer

***Relative_BrainWeight:*** Average of (Brain weight (mg)/ Body weight (g)) Encephalization coefficients.

***Longevity_Years_Estimate:*** Average lifespan of the species (in years) as estimated by the FishBase life history tool.

***Tmax:*** The maximum recorded age (years) of an individual of this species (recorded as the mean value across populations)

***Wmax:*** The maximum recorded weight (g) of an individual of this species (recorded as the mean value across populations)

***Lmax:*** The maximum recorded length (cm) of an individual of this species (recorded as the mean value across populations)

***TempPref_min/max and TempRange:*** Minimum and Maximum (and difference between) of the species preferred temperature range. This is often a generalization for the species and may not be representative of egg, larval, or juvenile stages.

***TolerableTemp_min/max and Temp_Range_Tolerable:*** Minimum, maximum and Tolerable temperature range of the species.
